# Selective granzyme B replenishment dominates the molecular switch for sustained cytotoxicity in effector CD8^+^ T lymphocytes

**DOI:** 10.64898/2026.10.09.758067

**Authors:** Hazyadee Frecia Rodriguez-Gutierrez, Anne-Laure Iscache, Aurèlie Montagne, Sabina Mueller, Salvatore Valitutti, Roxana Khazen

## Abstract

Cytotoxic T lymphocytes (CTLs) are highly efficient multiple killer cells, yet how they sustain killing after the initial discharge of canonical lytic granules remains incompletely understood. Late-phase cytotoxicity is often attributed to death receptor signaling or cytokine-mediated mechanisms, but whether CTLs also remodel their intrinsic cytotoxic machinery during prolonged engagement remains unclear. Here, we show that human CTLs maintain cytotoxic resilience through selective remodeling of their lytic program, marked by preferential preservation of granzyme B (GzmB). Following acute degranulation and reduction of mature perforin, CTLs selectively replenish GzmB through de novo protein synthesis, whereas perforin and several other granule-associated components show limited restoration. Replenished GzmB becomes redistributed into vesicular compartments with reduced overlap with mature perforin and is delivered to successive targets during sustained cytotoxic responses. Under conditions of limited detectable mature perforin, CTLs continue to deliver enzymatically active GzmB, induce altered calcium responses in target cells, and support productive but kinetically heterogeneous killing. This GzmB-preserved effector state is maintained and further accentuated during engagement with resistant melanoma cells. Together, our findings identify selective GzmB renewal as a central feature of sustained CTL cytotoxicity and reveal an adaptive remodeling of the cytotoxic machinery that may support CTL resilience during prolonged immune responses.

## Introduction

Cytotoxic T lymphocytes (CTLs) are central effectors of adaptive immunity, capable of eliminating virally infected and malignant cells with remarkable specificity^1^. Despite a high killing capacity, their presence within tumors does not necessarily translate into tumor control. In melanoma, for instance, spontaneously primed CD8⁺ T cells recognizing defined tumor antigens circulate in the blood, and metastatic lesions often contain abundant CTL infiltrates, yet disease progression and relapse remain common^2,3^. Immunotherapy has transformed cancer treatment by reviving exhausted T cells through checkpoint blockade, leading to durable responses in a subset of patients. However, the majority of patients either fail to respond or eventually relapse^4–6^. These observations suggest that the mere presence or reinvigoration of CTLs does not guarantee durable tumor clearance, rather the capacity of CTLs to sustain cytotoxic function over time (cytotoxic resilience) may be a critical limiting factor. How CTLs maintain killing competence under conditions of chronic stimulation and repeated target engagement therefore remains an important unresolved question.

Classically, CTLs are understood to rely on pre-formed cytotoxic granules containing perforin and granzymes that are rapidly released at the immunological synapse upon target recognition^1,7^. This framework explains the speed of the initial cytotoxic burst, but offers limited insight into how CTLs sustain killing capacity after early degranulation. Prior work has described shifts in target-cell death pathways, including FasL-mediated cytotoxicity in NK cells and regulated FasL mobilization in T cells under specific conditions^8,9^, yet these studies primarily address target-cell performance rather than CTL-intrinsic mechanisms that preserves cytotoxic resilience. Although post-degranulation recovery of lytic function has been documented, including through *de novo* synthesis and recycling of secretory compartments, it remains unclear whether CTLs replenish effector molecules in bulk, or instead undergo selective effector renewal. More broadly, how such regeneration shapes killing behavior during prolonged interactions with successive or resistant target cells remains unresolved. Because effective tumor control depends on repeated cytotoxic encounters, these questions raise the possibility that sustained engagement does not simply constrain cytotoxic output, but actively reshapes the molecular logic by which CTLs kill.

Here, we sought to determine how human CTLs preserve cytotoxic competence during sustained activation and repeated engagement with tumor targets. We show that human CTLs undergo a time dependent, effector-selective remodeling of their cytotoxic machinery during sustained activation. Following an initial perforin-and GzmB-dependent killing burst, CTLs rapidly deplete their lytic stores yet selectively replenish GzmB with minimal restoration of mature perforin. Using high dimensional flow cytometry, functional assays, live-cell imaging, and subcellular localization analyses, we demonstrated that newly synthesized GzmB is redistributed into noncanonical vesicular compartments and supports a late phase of cytotoxicity characterized by distinct target-cell calcium signatures. Importantly, this adaptive GzmB-centric program is preserved and amplified during CTL engagement with resistant melanoma targets, indicating that this effector-selective remodeling is maintained even under conditions of tumor-imposed stress. Together, our findings identify effector-selective renewal as a key feature supporting sustained CTL cytotoxicity and provide a framework for understanding how cytotoxic programs evolve during extended immune responses.

## Results

### Antigen stimulation triggers a conserved sequence of degranulation and selective GzmB replenishment

To define how CTLs deploy their cytotoxic machinery at the onset of target engagement, we performed a time-course analysis of lytic responses following antigenic stimulation of human CD8⁺ T cell clones conjugated with target cells presenting either a low concentration, 10 nM, or a saturating concentration, 10 µM, of cognate antigenic peptide.

Flow cytometry analysis revealed rapid CD3 internalization within minutes of conjugation, indicating robust and sustained TCR signaling that remained persistently reduced over the course of the experiment, with no detectable recovery during sustained target-cell contact at either peptide concentration (Fig. 1A). In parallel, CTLs showed marked surface exposure of CD107a/LAMP-1, which peaked within the first 120 minutes, indicating rapid engagement of CTL into lytic degranulation. CD107a surface expression declined sharply after approximately 4 hours of continuous contact with cognate targets under both low-dose and saturating antigenic stimulation (Fig. 1B-C). Analysis of both intracellular and surface CD107a pools revealed that although CD107a molecules were partially replenished intracellularly, they were not detectably re-exposed at the plasma membrane during prolonged stimulation. This suggests that later phases of CTL activation are unlikely to be maintained primarily through continued fusion of CD107a-positive canonical lytic granules (Supplementary Fig. 1).

**Figure 1.**
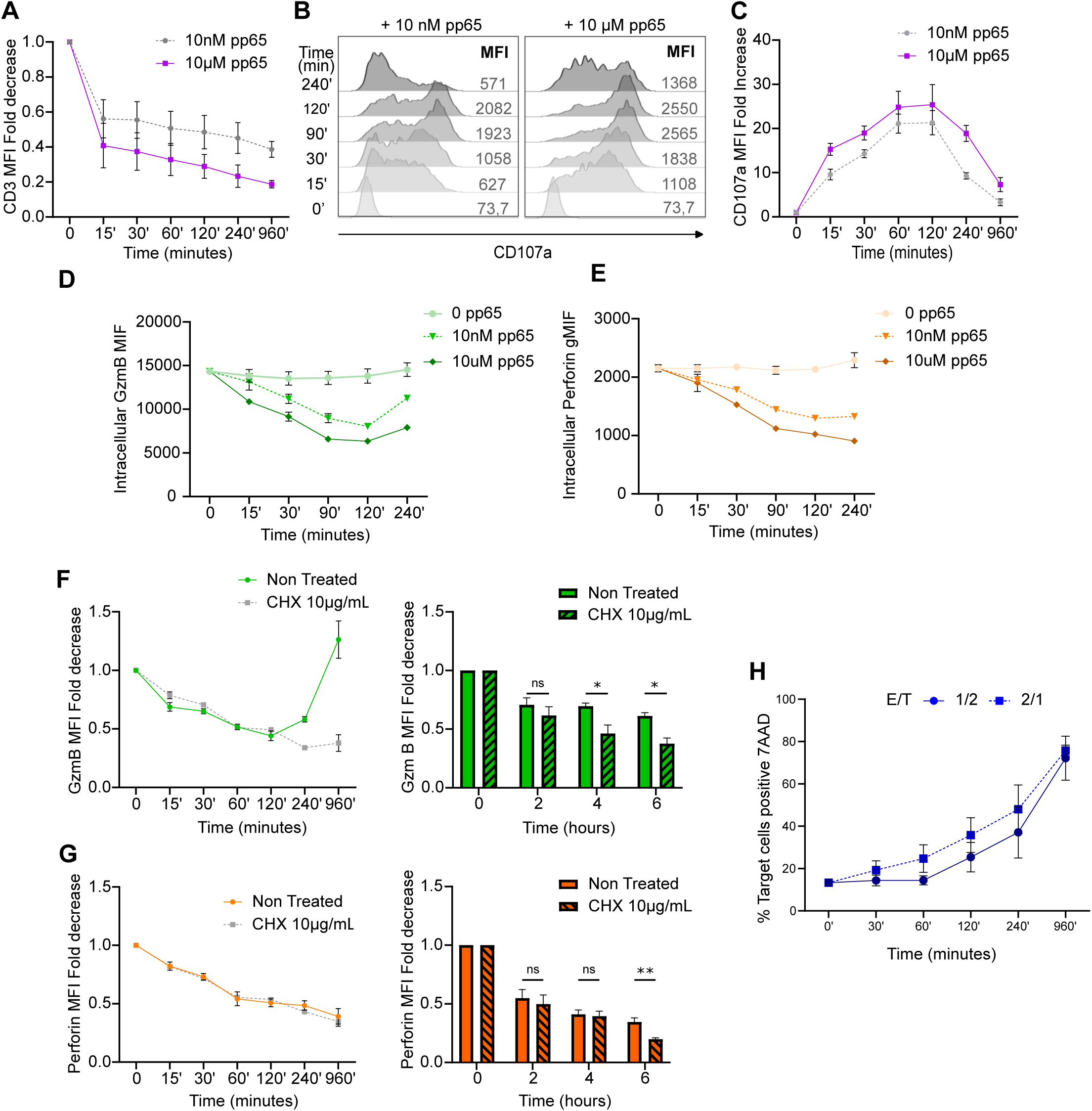
Antigen stimulation induces rapid lytic granule discharge followed by selective GzmB replenishment. Human CMV-specific CTLs were co-cultured with target cells either non-pulsed or pulsed with low (10 nM) or saturating (10 µM) concentration of cognate antigenic peptide for defined durations ranging from 15 minutes to 16 hours. **(A)** CD3 expression was monitored as an activation marker and internal control of CTL responsiveness to target cell recognition. CD3 expression was measured as mean fluorescence intensity (MFI) fold decrease relative to the NP condition. Pooled data from three independent experiments are shown. **(B-C)** Time-course analysis of CTL degranulation, measured as MFI fold increase of CD107a (LAMP1) surface exposure over NP condition. Representative histograms (B) and pooled quantification of CD107a MFI fold change relative to the NP condition (C) are shown. **(D-E)** Flow cytometric analysis of intracellular granzyme B (GzmB) (D) and perforin (E) levels in CTLs post conjugation with targets. Pooled MFI quantification from at least three independent experiments are shown. **(F-G)** Time kinetics (15min-16h) of normalized intracellular GzmB (F) and perforin (G) levels in CTLs conjugated with high-antigen targets in the presence or absence of cycloheximide (CHX; 10 µg ml⁻¹); right panels show summary quantification at the indicated time points. **(H)** Cytotoxicity was measured as the percentage of target cells positive for 7-AAD over time at indicated E:T ratios. Data are mean +/-s.e.m. from at least three independent experiments. Statistical significance was assessed by unpaired t-test. **ns, not significant; *P < 0.05; **P < 0.01.

In parallel, within this time frame, CTLs showed significant reduction in intracellular perforin and granzyme B (GzmB) content, consistent with robust granule discharge initiated as early as 15 minutes post target cell engagement (Fig. 1D-E). This early depletion was observed at both 10 nM and 10 µM peptide concentrations, indicating that low and saturating antigenic stimulation trigger a similar initial phase of cytotoxic effector deployment.

Importantly, kinetic analysis of GzmB expression revealed that, after this initial depletion phase, CTLs began to restore GzmB content by 4 hours post-conjugation, with levels progressively returning to baseline or exceeding baseline by 16 hours. In contrast, perforin expression remained low throughout this period (Fig. 1F-G). This divergent recovery pattern was detected at both antigen concentrations, suggesting that antigen dose modulates the strength of activation without altering the core sequence of effector remodeling: early granule discharge followed by selective GzmB restoration in the absence of comparable perforin recovery. To determine whether GzmB recovery resulted from de novo synthesis, CTLs were treated with cycloheximide (CHX) during target-cell engagement. CHX treatment completely abrogated GzmB replenishment, demonstrating that recovery depends on active protein synthesis rather than redistribution of residual intracellular stores (Fig. 1F-G).

To distinguish between total and functional perforin pools, perforin expression was assessed using antibodies recognizing either the mature, granule-associated form, dG9, or the total perforin pool, B-D48. While low levels of total perforin remained detectable, the mature perforin form was not restored following the initial degranulation phase, confirming that CTLs do not reconstitute the functional perforin pool with the same kinetics as GzmB during sustained activation (Supplementary Fig. 2).

The observation that selective GzmB resynthesis occurred not only at saturating antigen concentration but also at 10 nM peptide is particularly important, as this low-dose condition induces limited cytokine production as previously reported^10,11^. Thus, GzmB replenishment is unlikely to simply reflect a secondary consequence of maximal inflammatory signaling. Instead, it appears to represent a sensitive and robust feature of antigen-triggered CTL activation (Supplementary Fig. 3).

To relate early effector deployment to functional output, we measured target-cell death by quantifying 7-AAD uptake in target cells at matched time points (Fig. 1H). At different effector-to-target (E/T) ratios, including 1:2 and 2:1, target-cell death accumulated progressively over time rather than reaching an early plateau. This indicates that rapid granule discharge supports initial cytotoxicity, but that cumulative target-cell elimination during prolonged stimulation likely requires mechanisms beyond the initial pre-formed lytic granule pool.

Together, these data establish that acute CTL activation is characterized by rapid depletion of key lytic effectors, followed by selective GzmB resynthesis in the absence of comparable mature perforin restoration. These findings define a transient window of high granule-dependent cytotoxic output and reveal an early shift toward adaptive effector remodeling that may preserve cytotoxic resilience during continued target engagement.

### Sustained CTL activation selectively preserves GzmB while rewiring cytotoxic effector programs

To define how CTLs adapt their cytotoxic machinery following acute activation and degranulation, we performed high-dimensional spectral flow cytometry on human CTLs interacting with peptide-pulsed target cells under either low (10 nM) or high (10 µM) antigen stimulation. Because the initial kinetic analysis established that selective GzmB replenishment occurred under both low, 10 nM, and saturating, 10 µM, antigenic stimulation, subsequent phenotypic profiling was interpreted primarily under the saturating 10 µM condition to ensure robust and synchronized target-cell recognition. CTLs were analyzed after 2 hours, corresponding to the acute degranulation phase, 4hours as the time interval for GzmB recovery and 16 hours, corresponding to sustained target-cell engagement, using a 15-parameter panel encompassing major lytic effectors, degranulation markers, and complementary effector mediators (Fig. 2A).

**Figure 2.**
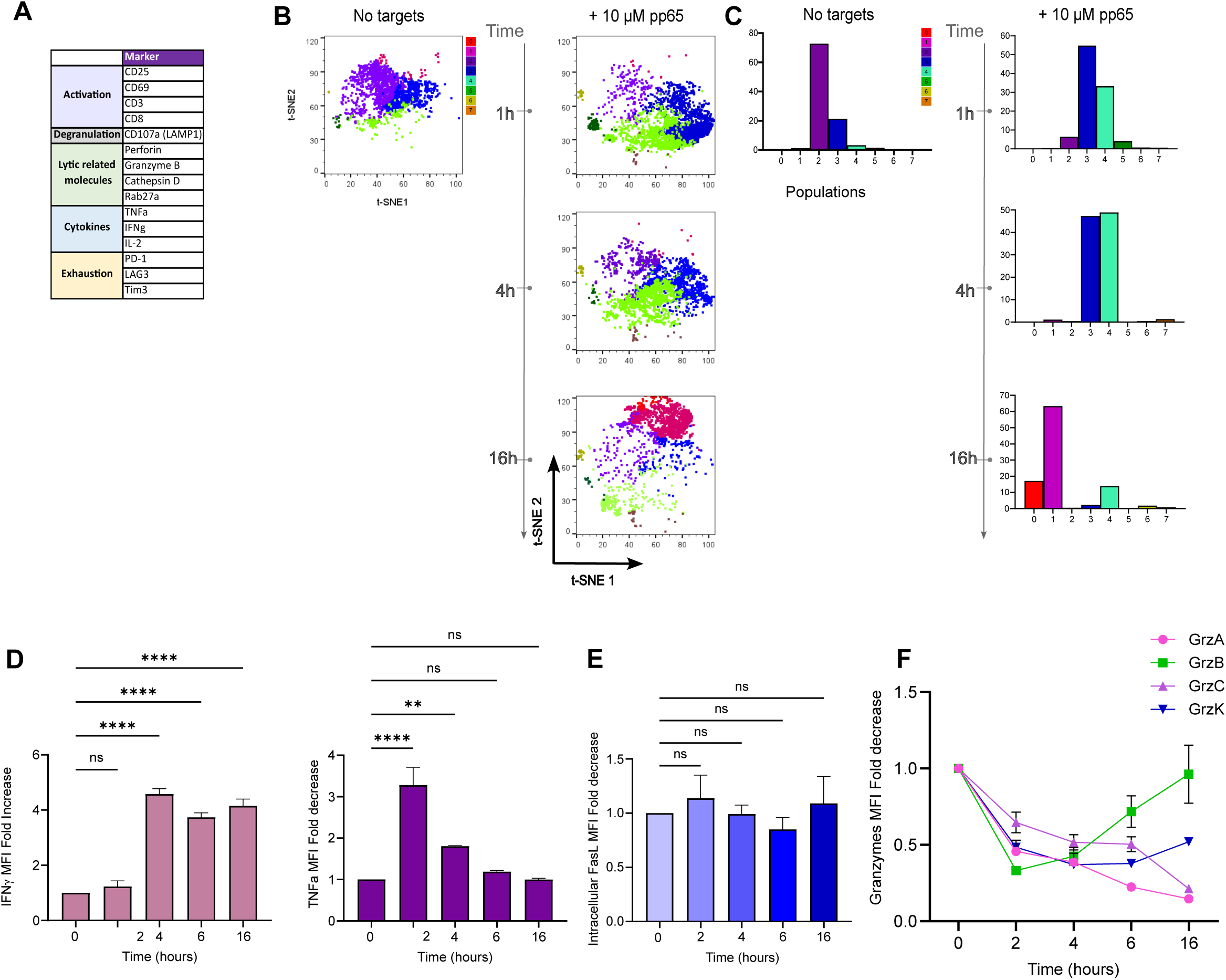
Sustained CTL activation is associated with selective preservation of GzmB and diversification of effector outputs. Human CTLs were conjugated with target cells either non-pulsed or pulsed with 10 µM cognate antigenic peptide for the indicated times and analyzed by flow cytometry. CTLs were gated as live CMFDA⁺ cells. **(A)** Antibody panel used for high-dimensional profiling. **(B)** Opt-SNE projection of CTL phenotypes across time following antigen-specific target-cell engagement. **(C)** FlowSOM clustering showing the distribution of phenotypic subsets across time (colors denote clusters). **(D)** Bar charts showing pooled kinetic quantification of cytokine expression in CTLs, including IFNy (left) and TNF (right) expressed as MFI fold change relative to the NP condition. **(E)** Pooled kinetic quantification of FasL expression in CTLs, expressed as MFI fold change relative to the NP condition. **(F)** Pooled quantification of granzyme expression in CTLs after target-cell engagement, expressed as fold change relative to the NP condition. Data are mean +/-s.e.m. from at least three independent experiments. Statistical significance was assessed by unpaired t-test. ns, not significant; *P < 0.05; **P < 0.01; ***P < 0.001; ****P < 0.0001.

Unbiased dimensionality reduction analysis revealed time-dependent changes in the CTL effector phenotype. Under saturating antigenic stimulation, CTLs analyzed after 2 hours and 16 hours occupied partially distinct regions of the Opt-SNE space, consistent with phenotypic remodeling during sustained target-cell engagement (Fig. 2B). This analysis was used as an exploratory visualization of global phenotypic changes rather than as a quantitative measure of antigen-dose-dependent differences. Consistently, phenotypic subsets detected by high-dimensional analysis showed changes in relative abundance and marker expression between the acute and sustained phases of activation (Fig. 2C). These data indicate that prolonged antigenic stimulation is accompanied by progressive remodeling of the CTL effector phenotype.

Kinetic evaluation of individual lytic molecules revealed that several components of the canonical lytic-granule machinery, including granzyme A, cathepsin D, and serglycin, declined following the acute degranulation phase and were not detectably restored during prolonged stimulation (Supplementary Fig. 4). In contrast, and consistent with the kinetics described above, GzmB expression was maintained or restored during sustained activation, distinguishing it from several other measured granule-associated components. This pattern occurred alongside induction of additional effector outputs, including Fas ligand (FasL) expression and production of inflammatory cytokines such as IFNγ and TNFα (Fig. 2D-E). Thus, sustained CTL activation was associated with selective preservation of GzmB together with increased expression of non-granule-associated effector mediators, rather than with uniform restoration of the initial lytic-granule content (Fig. 2E).

Together, these data show that, following acute degranulation, human CTLs undergo measurable changes in their cytotoxic effector profile during sustained target-cell engagement. Rather than restoring all measured canonical lytic-granule components with similar kinetics, CTLs selectively restore GzmB, while perforin and several additional granule-associated molecules remain reduced. In parallel, CTLs increase expression of complementary effector mediators, including FasL and inflammatory cytokines. These findings suggest that sustained CTL activation is accompanied by a shift in effector composition, in which GzmB preservation occurs alongside broader changes in cytotoxic and inflammatory output.

### Sustained CTL activation spatially uncouples GzmB from perforin-containing lytic granules

To investigate where replenished GzmB localizes during prolonged cytotoxic activity, we examined the subcellular distribution of GzmB and perforin by confocal imaging in CTLs conjugated with peptide-pulsed target cells for 0, 4, or 16 hours (Fig. 3A). Before interaction with target cells (Time 0), GzmB and perforin displayed overlapping punctate staining patterns within CTLs, consistent with their localization in intracellular vesicular structures. However, at later time points (4-16 hours), GzmB staining remained readily detectable, whereas its intracellular distribution became progressively altered. Specifically, GzmB was observed in vesicular structures that were spatially distinct from perforin-positive compartments (Fig. 3A). Quantification of voxel overlap confirmed this spatial segregation. The proportion of GzmB signal co-localizing with perforin, GzmB/Perf overlap, decreased significantly over time (Fig. 3B), whereas the remaining perforin-positive structures largely retained overlap with GzmB. These results suggest that, following initial degranulation, a growing fraction of GzmB accumulates in vesicular compartments with little or no detectable mature perforin, consistent with progressive spatial uncoupling of these two lytic effectors over time.

**Figure 3.**
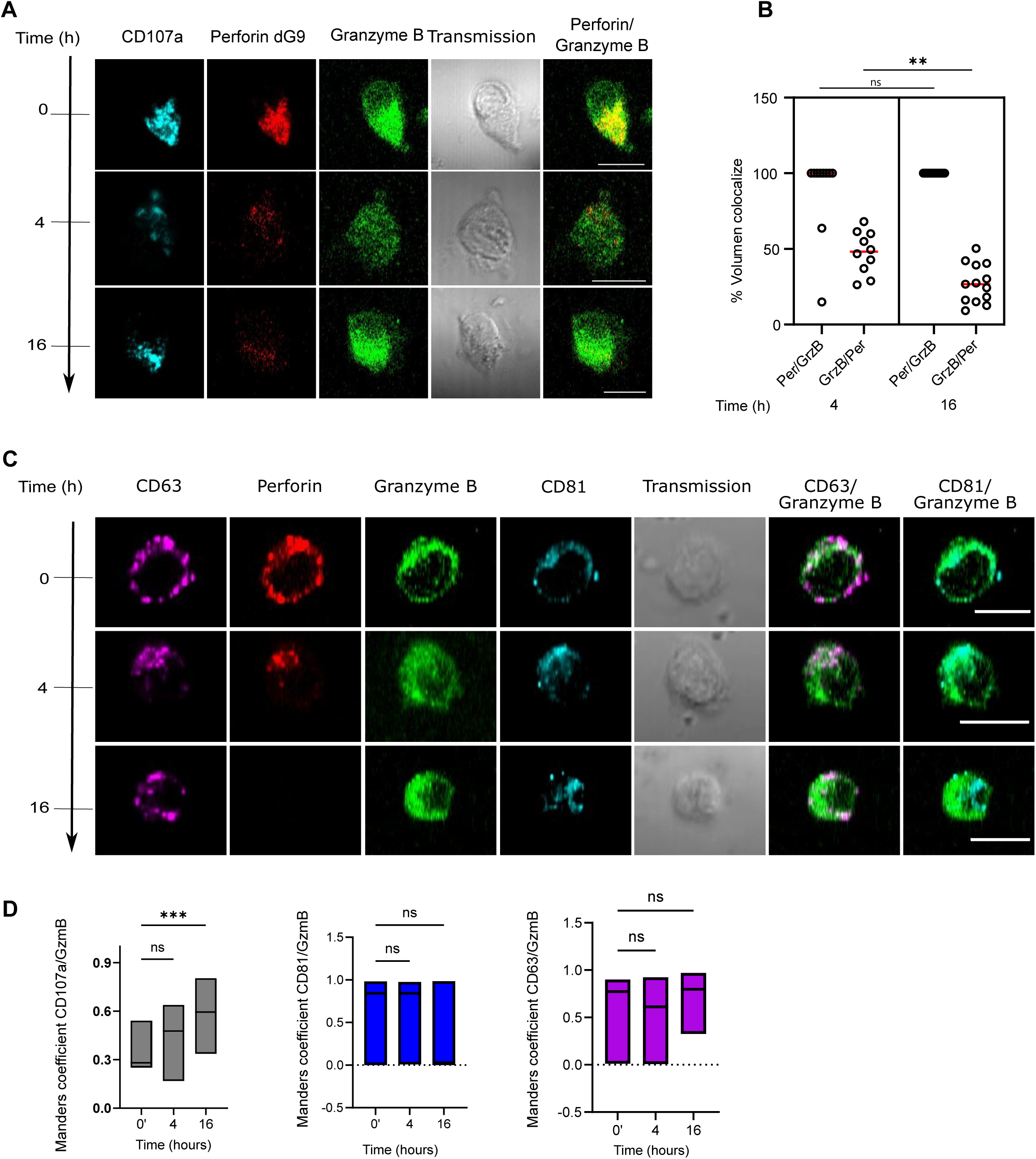
Sustained CTL activation uncouples granzyme B from perforin and redistributes it into noncanonical vesicular compartments. **(A)** Representative confocal images of CTLs conjugated with peptide-pulsed target cells for 0, 4, or 16 h, showing CD107a (LAMP1, Cyan), GzmB (green) and perforin (red). **(B)** Quantification of voxel-level colocalization, shown as the fraction of GzmB signal overlapping with perforin (GzmB/Perf) and the fraction of perforin signal overlapping with GzmB (Perf/GzmB) at the indicated time points. **(C)** Representative confocal images showing GzmB (green) and perforin (red) relative to CD63 (purple) and CD81 (cyan) at the indicated time points. **(D)** Quantification of GzmB association with vesicular markers, expressed as Pearson’s correlation coefficient between GzmB and CD107a, CD63, or CD81 signals at the indicated time points. Data are presented as mean ± s.e.m. Statistical significance was assessed using unpaired t-test. ***P < 0.001; ****P < 0.0001.

To further characterize the vesicular compartments associated with GzmB during sustained cytotoxic engagement, we examined the localization of GzmB relative to the tetraspanins CD107a, CD63, and CD81, which mark partially overlapping endo-lysosomal, secretory, and recycling-associated vesicular populations. Confocal imaging revealed that CD107a-, CD63-and CD81-positive vesicular structures were maintained throughout prolonged interaction with target cells (Fig 3A and C). To determine whether the persistence of these vesicular compartments reflected active turnover rather than passive retention, we analyzed CD63 and CD81 expression by flow cytometry in the presence or absence of CHX. Inhibition of protein translation significantly reduced the expression of both CD63 and CD81 during prolonged activation, consistent with the CHX-sensitive maintenance of the intracellular CD107a pool (Supplementary Fig. 5), indicating the steady-state expression of these vesicular markers during prolonged activation is maintained through ongoing de novo synthesis rather than passive retention.

Within this dynamically maintained vesicular pool, quantitative co-localization analysis revealed that while GzmB was detected within multiple vesicular types, it displayed the highest degree of partial spatial overlap with CD107a⁺ compartments during the sustained phase of cytotoxicity (Fig. 3D). This heterogeneous redistribution of GzmB across vesicular subsets during repetitive target encounter, suggests that CTLs deploy distinct intracellular routing strategies to maintain access to this key cytotoxic effector.

Together, these data demonstrate that sustained CTL activation induces a pronounced spatial reorganization of the cytotoxic machinery, whereby newly synthesized GzmB becomes uncoupled from perforin-containing lytic granules and redistributed into non-canonical vesicular compartments. This architectural remodeling provides a mechanistic framework for how CTLs preserve GzmB availability despite sustained perforin depletion, thereby supporting alternative cytotoxic strategies during prolonged target engagement.

### GzmB renewal and redeployment support CTL cytotoxic resilience during sequential target challenges

To determine whether GzmB replenishment is associated with sustained cytotoxic function, we assessed cumulative target-cell killing over time by CTLs conjugated with peptide-pulsed target cells at different E:T ratios, ranging from 1:20 to 2:1. Cytotoxicity was quantified by 7-AAD uptake in target cells. Across all E:T ratios, cumulative target-cell death increased between 4 and 16 hours, indicating that CTLs retain cytotoxic activity during prolonged target-cell exposure (Fig. 4A). Notably, continued killing was still observed under the lowest E:T conditions, where individual CTLs are expected to encounter multiple targets and where the contribution of collective effector activity is minimized. Although this assay does not directly track individual CTLs, these findings are consistent with preserved cytotoxic competence under conditions that favor repeated target engagement by single effector cells.

**Figure 4.**
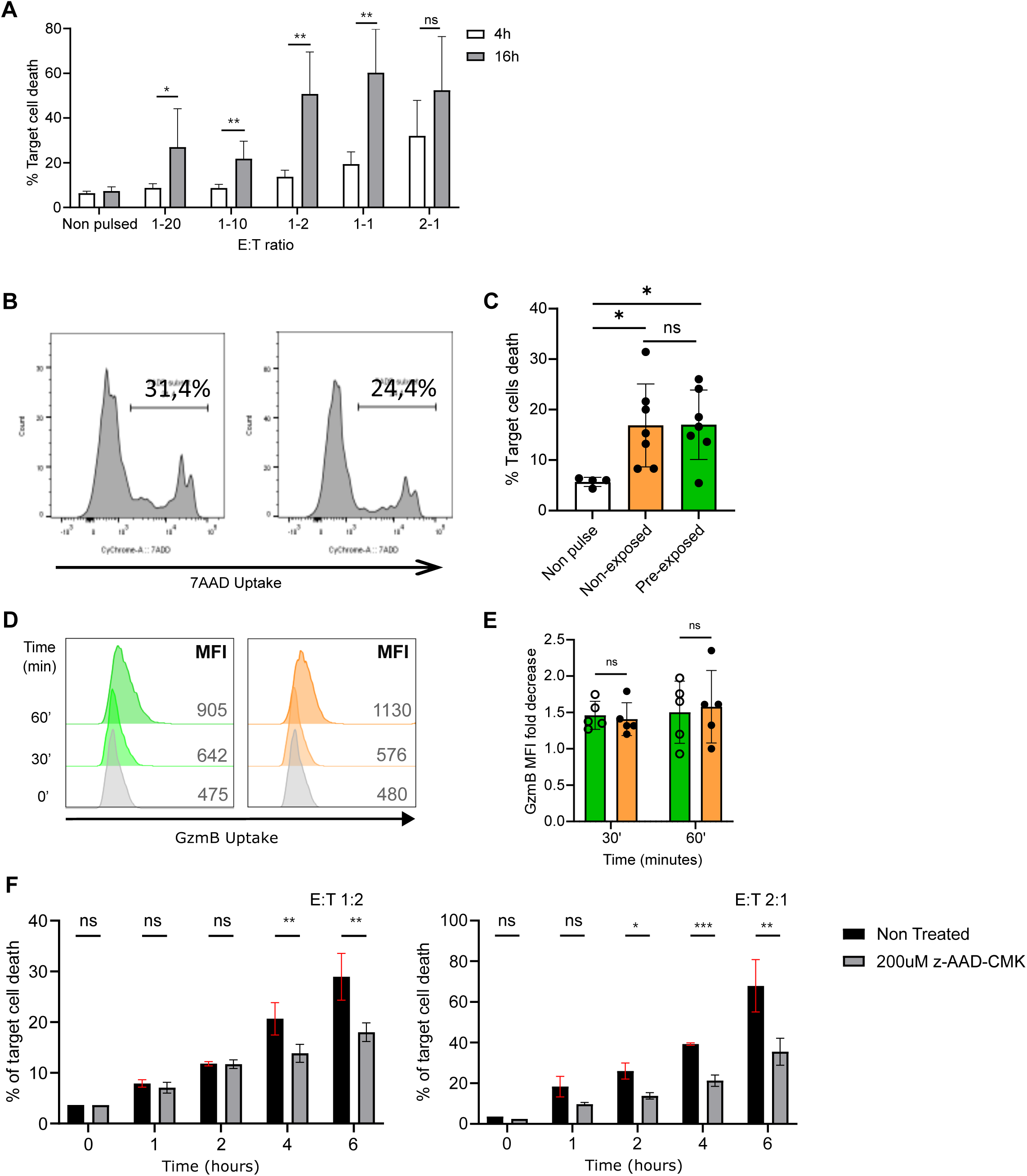
Granzyme B replenishment supports sustained cytotoxicity during continuous and serial target-cell challenge. **(A)** CTL-mediated cytotoxicity measured at 4 h and 16 h across increaseing E:T ratios, quantified as the percentage of 7-AAD⁺ target cells. **(B-C)** CTLs were first conjugated with peptide-pulsed target cells (10 µM) for 4 h to induce degranulation and perforin depletion, then conjugates were mechanically dissociated and “pre-exposed” CTLs were re-challenged with a second batch of fresh peptide-pulsed targets. B) Representative flow cytometry histograms of 7-AAD staining in target cells conjugated with fresh or pre-exposed CTLs. C) Pooled quantification of cytotoxicity from three independent experiments. **(D)** Representative histograms showing GzmB uptake in target cells following conjugation with fresh or pre-exposed CTLs. **(E)** Pooled quantification of target-cell GzmB uptake. **(F)** CTLs were conjugated with target cells unpulsed or pulsed with cognate antigenic peptide in the presence or absence of granzyme B inhibitor. Cytotoxicity was evaluated as the percentage of 7-AAD⁺ target cells. Two E:T ratio were tested as indicated. Data are shown as mean +/-s.e.m. Statistical significance was assessed using unpaired t-test. ns, not significant, *P < 0.05; **P < 0.01; ***P < 0.001; ****P < 0.0001.

To directly test whether cytotoxicity is sustained across sequential encounters, we implemented a two-phase killing assay. CTLs were first conjugated with peptide-pulsed target cells for 4 hours to induce degranulation and reduce intracellular perforin. These pre-exposed CTLs were then re-challenged with a fresh batch of cognate targets (Fig. 4B). Despite prior activation and lytic granule discharge, pre-exposed CTLs retained similar cytotoxic activity against newly added targets (Fig. 4B-C).

To determine whether this second wave of killing was associated with renewed GzmB deployment, we measured GzmB transfer to peptide-pulsed target cells in the two-phase assay (Fig. 4D-E). GzmB was detected in target cells conjugated to both non-exposed and pre-exposed CTLs, indicating that previously challenged CTLs remain capable of delivering GzmB during secondary target engagement. Because sustained activation was also associated with induction of complementary effector pathways, including cytokine production and FasL expression, we next assessed whether these pathways contributed to late cytotoxic activity. Blockade of calcineurin-dependent signaling, which is known to inhibit cytokine expression in T cells^12^, or inhibition of FasL did not significantly impair late cytotoxicity (Supplementary Fig. 6A-B). In contrast, pharmacological inhibition of GzmB markedly reduced sustained killing (Fig. 4F). These findings indicate that, under these experimental conditions, late cytotoxic activity depends primarily on GzmB activity rather than on calcineurin-dependent cytokine responses or FasL-mediated cytotoxicity.

Together, these results indicate that CTLs preserve cytotoxic competence during repeated target challenges through selective GzmB replenishment and redeployment, despite limited restoration of detectable mature perforin. Thus, cytotoxic resilience in human CTLs is supported by a GzmB-centric adaptive program that enables sustained killing during prolonged immune responses when the canonical perforin-containing granule pool is not fully restored.

### Sustained CTL engagement shifts target-cell Ca²⁺ responses away from rapid perforin-like spiking

To determine how CTLs execute sustained target-cell cytotoxicity under conditions of reduced mature perforin availability, we examined intracellular calcium dynamics in target cells using live-cell imaging. The experimental design paralleled the sequential killing assay from Figure 4B. Specifically, CTLs were first conjugated with peptide-pulsed target cells for 4 hours to induce degranulation and perforin reduction. These pre-exposed CTLs were then re-challenged with fresh peptide-pulsed target cells under live-imaging conditions in the presence of the fluorescent calcium reporter Fluo-4.

During early phase of cytotoxicity, attacks mediated by fresh CTLs, 0-2 hours, induced characteristic sharp and repetitive intracellular Ca²⁺ spikes in target cells (Fig. 5A, C), consistent with rapid membrane perturbation and pore-forming activity classically associated with perforin-dependent cytotoxicity. In contrast, target cells engaged by pre-exposed CTLs, corresponding to the 4-6-hour phase of sustained engagement, displayed qualitatively altered Ca²⁺ profiles. These included faint single transient increases or slow Ca²⁺ accumulation, with little or no repetitive spiking behavior (Fig. 5B, C). Across analyzed target cells, these responses were of markedly lower amplitude, indicating a substantial shift in the dynamics of target-cell Ca²⁺ entry during late CTL engagement.

**Figure 5.**
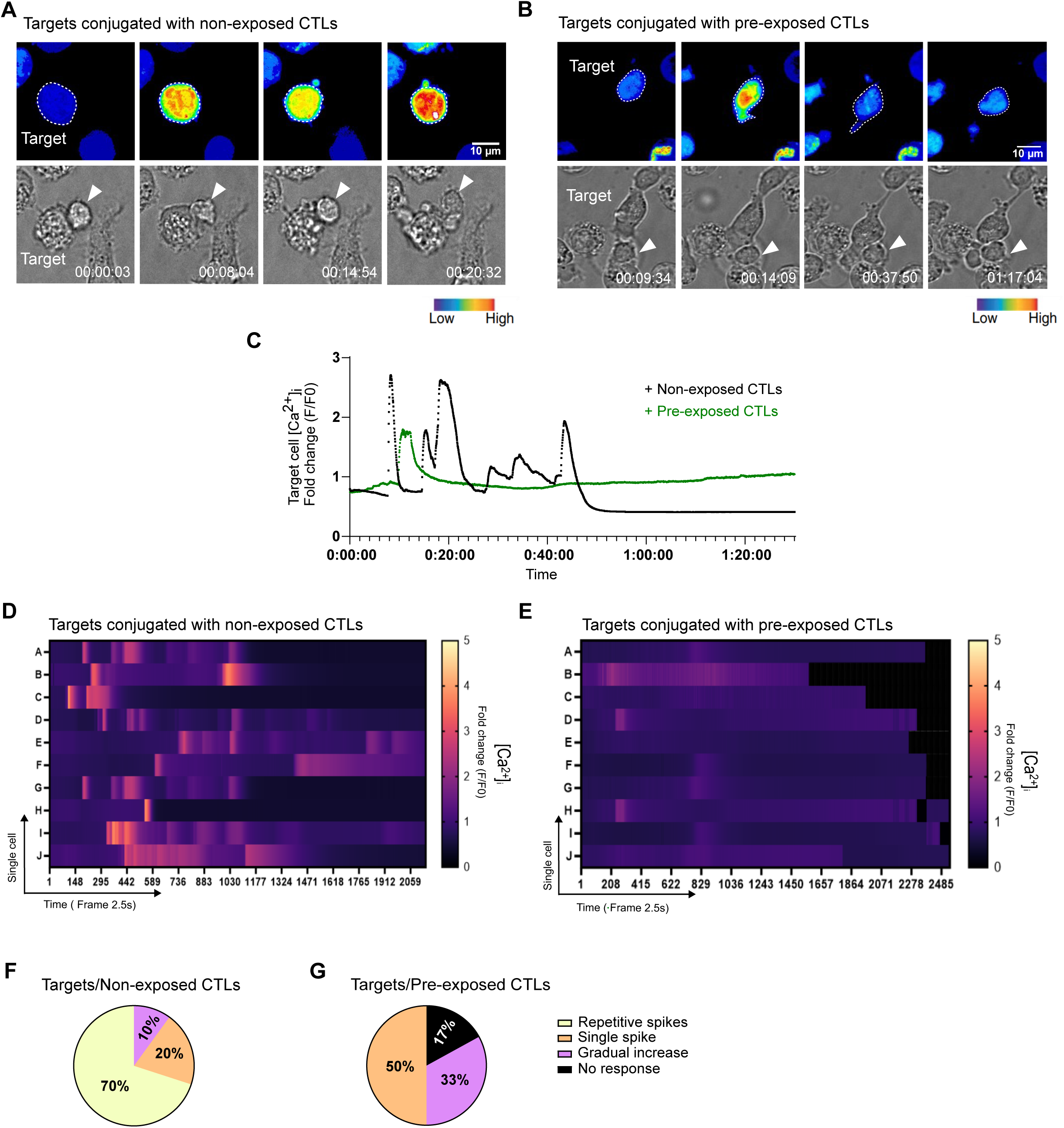
Sustained CTL cytotoxicity under perforin depletion is associated with distinct target-cell Ca²⁺ signatures. CTLs were first conjugated with peptide-pulsed target cells to induce degranulation and perforin depletion for 4h (pre-exposed CTLs), then rechallenged with a second batch of fresh peptide-pulsed targets preloaded with Fluo-4 to monitor intracellular calcium dynamics. (fresh/non-exposed CTLs: acute phase behavior, Pre-exposed CTLs: sustained phase behavior) **(A)** Representative time-lapse snapshots of intracellular Ca²⁺ levels ([Ca²⁺]ᵢ, pseudocolor) in target cells during killing by fresh CTLs (early phase, 0-2 h). White dashed line outlines the target cell. **(B)** Representative snapshots as in (A) during killing by pre-exposed CTLs (sustained phase, 4-6 h), highlighting altered target-cell Ca²⁺ dynamics. **(C)** Representative single-cell [Ca²⁺]ᵢ traces from target cells conjugated with a fresh CTL (black) or a pre-exposed CTL (green) during the indicated phases. **(D,E)** Single-cell heatmaps of target-cell [Ca²⁺]ᵢ dynamics, displayed as pseudocolor intensity over time; each row represents an individual target cell. Traces are aligned to the first stable CTL/target contact. **(F,G)** Classification of target-cell Ca²⁺ responses into repetitive spikes, gradual increases, or no response, with category frequencies shown as percentages (pie charts) n = 20. White arrowheads indicate T cells. Scale bars, 10 µm.

Single-cell heatmaps and categorical analysis of Ca²⁺ response types revealed a clear temporal shift in target-cell signaling. Whereas most targets engaged by fresh CTLs exhibited rapid, repetitive Ca²⁺ spikes, those interacting with pre-exposed CTLs more frequently displayed gradual increases, single transient responses, or minimal detectable Ca²⁺ activity (Fig. 5D-G). Given the markedly reduced abundance of mature perforin in pre-exposed CTLs, together with the qualitative shift in target-cell Ca²⁺ dynamics, these data suggest a change in the mode of target-cell membrane perturbation during the late phase of cytotoxicity.

Together, these target-cell Ca²⁺ signatures indicate a time-dependent shift in how CTLs perturb target cells during sustained engagement, consistent with a reduced contribution of rapid perforin-mediated membrane permeabilization during late cytotoxic responses.

### GzmB-mediated cytotoxic resilience preserves killing with heterogeneous functional kinetics

To determine whether the altered Ca²⁺ signatures observed during sustained cytotoxicity correspond to effective target-cell killing, we directly assessed GzmB delivery and enzymatic activity in target cells. Using both flow cytometry and live-cell imaging, we monitored cleavage of a fluorogenic GzmB substrate, GranToxiLux, during CTL-target interactions. CTLs were first pre-exposed to peptide-pulsed targets to induce degranulation and reduce detectable mature perforin, then re-challenged with fresh targets under conditions identical to the experimental conditions described above.

Live-cell video microscopy of conjugates formed by either fresh, non-exposed CTLs, or pre-exposed CTLs with peptide-pulsed target cells revealed clear cleavage of the GzmB substrate in target cells, indicating delivery of enzymatically active GzmB during sustained cytotoxic responses (Fig. 6A). Consistently, flow cytometric analysis confirmed that GzmB substrate cleavage was preserved in targets conjugated with pre-exposed CTLs compared with those conjugated with fresh CTLs (Fig. 6B), indicating that functional GzmB activity in target cells is maintained during sustained cytotoxic engagement. Importantly, this activity was maintained despite a marked reduction in detectable mature perforin, suggesting that productive GzmB-mediated cytotoxicity can persist under conditions in which the mature perforin pool is strongly reduced (Fig. 1D-E).

**Figure 6.**
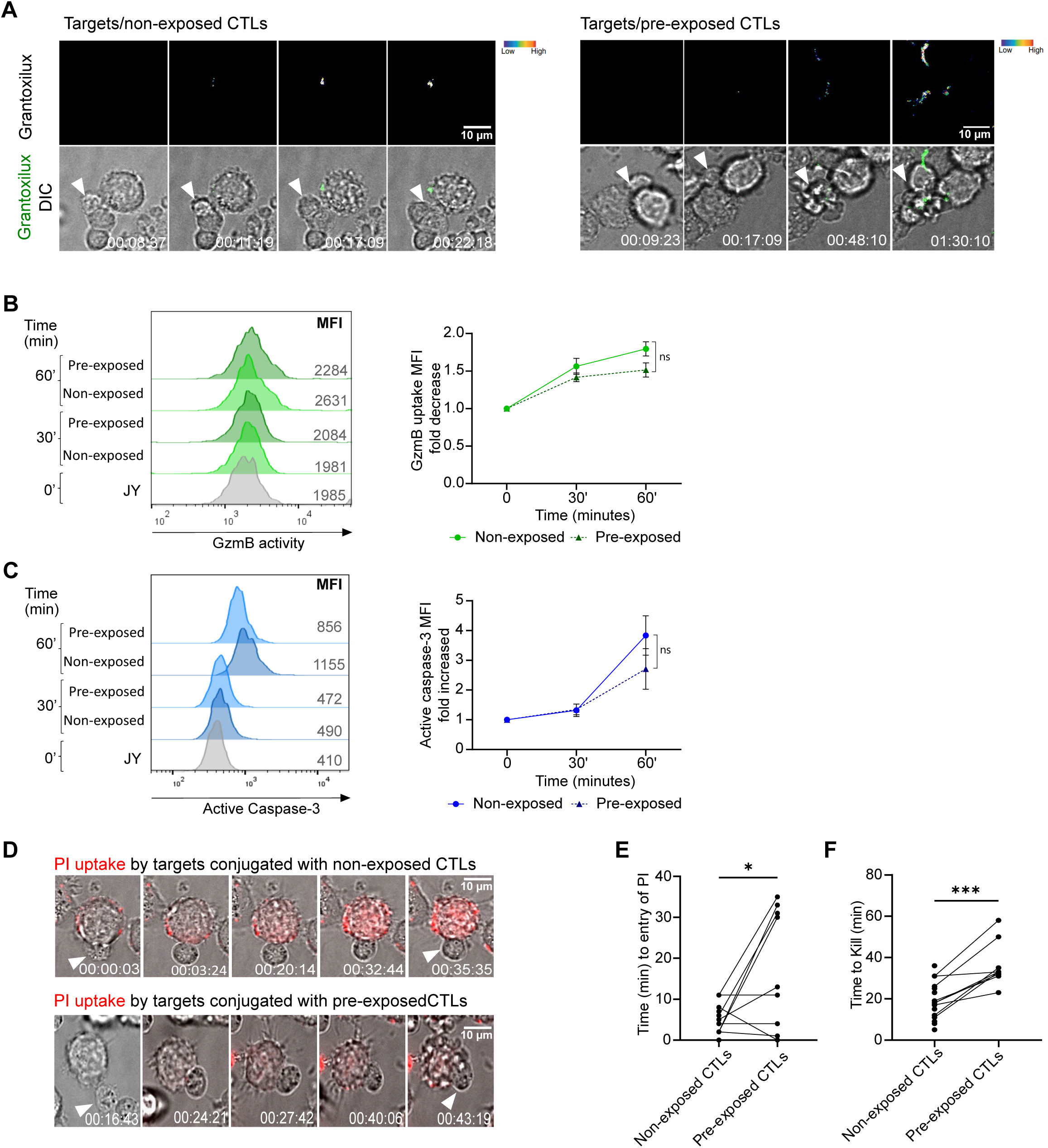
Enzymatically active granzyme B is delivered during sustained cytotoxicity and is associated with heterogeneous killing kinetics. Target cells were preloaded with a fluorogenic GzmB substrate (GranToxiLux) and conjugated with fresh or pre-exposed CTLs. **(A)** Representative time-lapse images showing GranToxiLux cleavage in target cells (green and pseudocolor) following interaction with fresh or pre-exposed CTLs; **(B)** Flow cytometric quantification of GranToxiLux cleavage in target cells at 30 min and 1 h after conjugation. Left, representative histogram; right, pooled quantification from n = 3 independent experiments. **(C)** Representative flow cytometry quantification of cleaved caspase-3 in target cells following conjugation with CTLs (same experimental conditions as in B). Left, representative histogram; right, pooled quantification from at least three independent experiments. **(D)** Representative snapshots from live-cell imaging showing PI uptake (red) in target cells as a marker of membrane permeabilization and cell death. Top, non-exposed/fresh (NE) CTLs; bottom, pre-exposed (PE) CTLs. White arrowheads indicate T cells. **(E)** Time to PI entry for target cells killed by non-exposed/fresh (NE) or pre-exposed (PE) CTLs. Data are shown as mean +/-s.e.m. Statistical significance was assessed using unpaired t-test. ns, not significant, *P < 0.05; ***P < 0.001.

To determine whether GzmB substrate cleavage was associated with productive target-cell death rather than abortive enzyme transfer, we assessed caspase-3 activation in target cells in parallel. Target cells engaged by either fresh or pre-exposed CTLs exhibited detectable GzmB substrate cleavage together with caspase-3 activation within 1 hour of target-cell engagement (Fig. 6C), confirming that sustained CTL engagement supports functional, cell death-inducing GzmB delivery.

To evaluate whether this time-dependent rewiring of the cytotoxic machinery alters the kinetics of target-cell killing, we performed live-cell imaging to monitor CTL-target interactions in the presence of propidium iodide, PI, as a marker of membrane permeabilization and cell death. Imaging confirmed that both fresh and pre-exposed CTLs induced PI uptake in target cells (Fig. 6D-E). However, whereas fresh CTLs typically induced rapid PI entry shortly after target-cell engagement, pre-exposed CTLs exhibited delayed PI entry and greater kinetic variability (Fig. 6E). Specifically, PI entry times were distributed across both faster and slower killing events, revealing increased heterogeneity in cytotoxic kinetics during sustained engagement. These data indicate that cytotoxic resilience is maintained during sustained engagement but is accompanied by a broader distribution of functional killing kinetics at the single-cell level.

Consistent with this kinetic heterogeneity, the overall time required for target-cell death was increased for pre-exposed CTLs compared with fresh CTLs (Fig. 6D, E). These findings indicate that while CTLs retain the capacity for productive GzmB-associated killing after prior degranulation, the tempo of cytotoxic performance becomes more heterogeneous and, on average, slower during sustained engagement.

Notably, this functional heterogeneity parallels the broadened distribution of GzmB expression observed after prolonged activation, at 16 hours, in which CTLs displayed a wider range of intracellular GzmB levels compared with the more homogeneous baseline state (Supplementary Fig. 7). Together, these data are consistent with a model in which the broadened distribution of intracellular GzmB levels that emerges during sustained activation is associated with increased intercellular variability in killing kinetics.

Together, these results establish that sustained CTL cytotoxicity proceeds through preserved delivery of enzymatically active GzmB, but with altered and heterogeneous killing kinetics. This variability represents a functional feature of the GzmB-centric resilience program and may help explain the heterogeneous cytotoxic behaviors that emerge during prolonged immune responses.

### Resistant melanoma targets preserve and amplify the GzmB-centric cytotoxic program

To determine whether the cytotoxic resilience program defined above is maintained during engagement with resistant tumor targets, we examined CTL responses during prolonged interaction with melanoma target cells. Although the melanoma cells used in this assay display reduced susceptibility to CTL-mediated killing, we have previously shown that CTLs can retain measurable cytotoxic activity upon antigen recognition in this setting^13–15^. Consistent with this, CTLs engaging melanoma cells displayed sustained cytotoxic activity over prolonged interaction, indicating that resistant targets do not fully abrogate CTL function but instead impose a context of prolonged cytotoxic stress.

To assess whether engagement with resistant targets induces a distinct or altered effector state, we next performed functional phenotyping of CTLs following conjugation with melanoma or EBV-transformed target cells. Dimensionality reduction analysis using Opt-SNE revealed that, across antigen doses and time points, 1, 4, and 16 hours, CTLs interacting with melanoma targets underwent a coordinated phenotypic displacement that followed a trajectory comparable to that observed during prolonged interaction with sensitive targets (Fig. 7A). Moreover, unsupervised clustering revealed the emergence and resolution of comparable phenotypic subsets over time in both contexts (Fig. 7B), indicating that the global effector reprogramming associated with sustained cytotoxic engagement is conserved across target types and is not restricted to highly permissive killing conditions.

**Figure 7.**
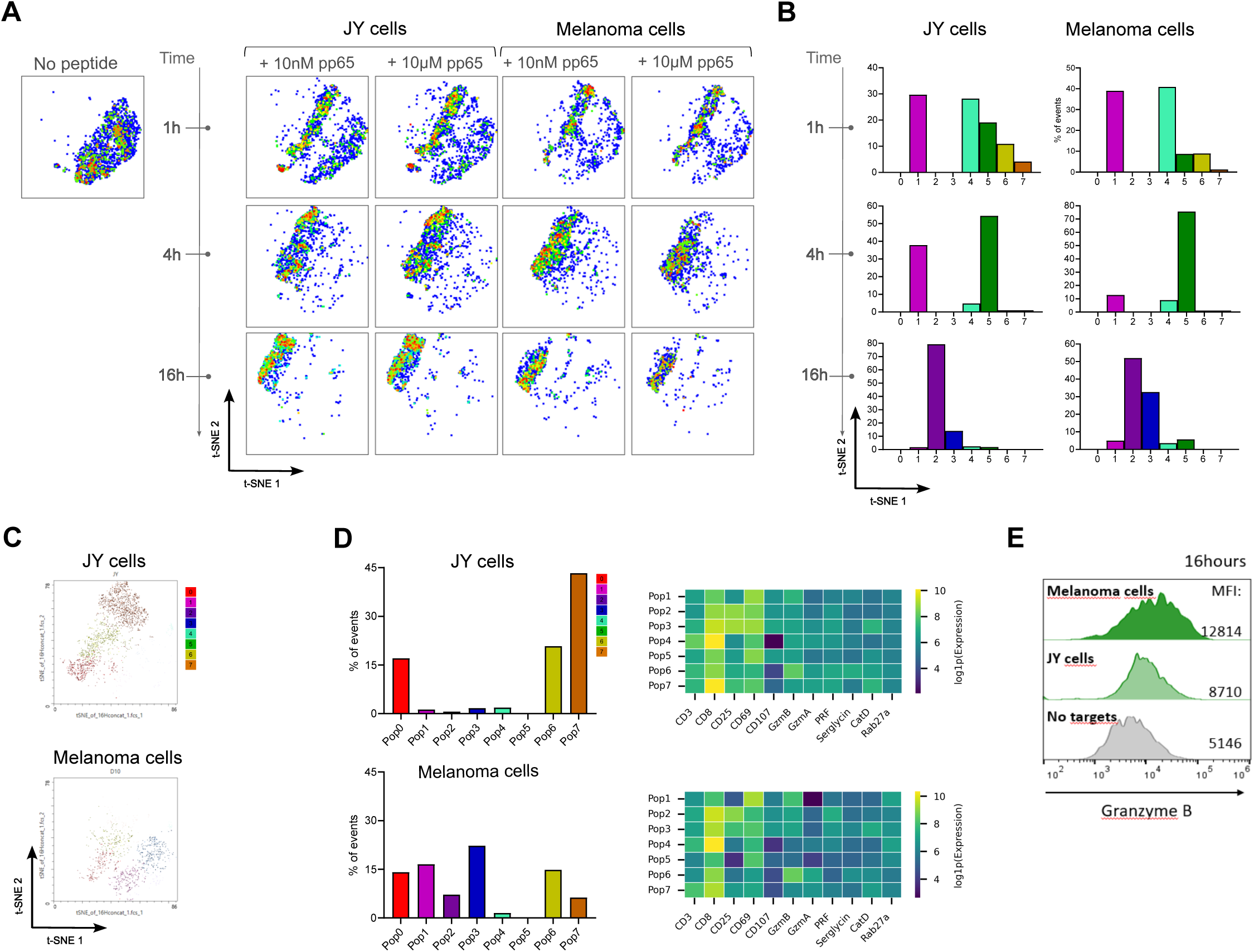
**Cytotoxic reprogramming is conserved during engagement with resistant melanoma targets and is associated with increased granzyme B abundance and heterogeneity**. CTLs were conjugated with sensitive (JY cells) or resistant melanoma target cells pulsed with cognate peptide at low or high antigen concentrations for the indicated times (1, 4, and 16 h) and analyzed by high-dimensional spectral flow cytometry. CTLs were gated as live CMFDA⁺ cells. **(A)** Opt-SNE projection of CTL phenotypes across target type (JY versus melanoma), antigen dose (low versus high), and time, showing comparable time-dependent phenotypic displacement under both target conditions. **(B)** FlowSOM-based clustering showing the distribution of phenotypic subsets over time for CTLs conjugated with JY or melanoma targets (high-antigen condition shown. **(C)** Opt-SNE projection restricted to the sustained phase (16 h) comparing CTLs conjugated with high-antigen JY or melanoma targets. **(D)** FlowSOM subset assignment overlaid on the 16 h Opt-SNE map, revealing melanoma-associated enrichment of subsets characterized by reduced perforin and increased heterogeneity in GzmB expression. **(E)** Representative histograms of intracellular GzmB in CTLs at 16 h comparing CTLs conjugated with high-antigen JY, melanoma targets, or CTLs cultured alone. Data are shown as mean +/-s.e.m. Statistical significance was assessed using unpaired t-test.

We next focused on CTLs undergoing prolonged engagement, 16 hours, to determine whether resistant targets modulate the composition or intensity of this adaptive state. Opt-SNE analysis of all conditions indicated that CTLs conjugated with melanoma targets occupied phenotypic regions overlapping with, yet distinct from, those engaged with the sensitive JY targets (Fig. 7A). When the analysis was restricted to the sustained phase, 16 hours, CTLs conjugated with melanoma cells segregated into distinct subsets characterized by reduced perforin expression and increased heterogeneity in GzmB levels (Fig. 7C-D). Consistently, CTLs interacting with melanoma cells exhibited significantly elevated and more broadly distributed GzmB expression compared with CTLs engaging sensitive targets or CTLs cultured alone (Fig. 7E). This amplification of GzmB abundance and heterogeneity occurred despite persistently low levels of detectable mature perforin and continued reduction of canonical lytic-granule features, suggesting that resistant tumor targets further reinforce the GzmB-centric effector program associated with cytotoxic resilience.

Together, these results indicate that CTL engagement with resistant melanoma targets preserves the adaptive cytotoxic reprogramming observed during prolonged interactions with sensitive cells while further accentuating GzmB abundance and heterogeneity.

## Discussion

CTLs are capable of executing successive rounds of target-cell killing, a property central to their effectiveness in antiviral and antitumor immunity ^16–20^. Yet most mechanistic understanding of CTL cytotoxicity derives from the first minutes of target recognition, when preformed lytic granules are rapidly mobilized toward the immunological synapse^1,10,15,21^. Serial killing can initially be sustained through economical use of pre-existing stores: CTLs release only a fraction of their available lytic granules during individual killing events, thereby preserving a reserve for subsequent targets^22,23^. Prolonged cytotoxicity, however, additionally requires re-arming of the effector machinery through membrane recycling and renewed biosynthesis. What happens to the cytotoxic machinery once this preformed arsenal has been deployed is considerably less understood. Here, we show that sustained cytotoxicity is not accompanied by equivalent restoration of the original lytic program. Instead, CTLs rebuild their effector machinery selectively. GzmB is actively replenished through *de novo* synthesis and becomes increasingly prominent, whereas detectable mature perforin and several other granule-associated components remain reduced. Importantly, this remodeled state remains functionally competent, allowing previously engaged CTLs to deliver enzymatically active GzmB and kill newly encountered targets, although with altered and increasingly heterogeneous kinetics. These findings therefore support a model in which prolonged CTL activity involves not simply replenishment of exhausted cytotoxic stores, but a time-dependent reconfiguration of the machinery through which killing is executed, in which GzmB renewal becomes a functional bottleneck for continued killing.

That CTLs can synthesize new cytotoxic proteins while responding to antigen is not itself new. Early studies established that TCR stimulation couples degranulation to renewed production of lytic proteins and identified secretion routes that are not restricted to preformed granules^24^. More recent work has further shown that cytotoxic re-arming is an actively supported biosynthetic process: inhibition of mitochondrial translation impairs replenishment of perforin and GzmB while leaving CTL motility and secretory competence comparatively preserved^25^. Newly synthesized perforin was shown to reach the immunological synapse rapidly and, under some conditions, to traffic through pathways partially distinct from conventional cytotoxic granules^26^. Our findings extend these observations by showing that renewal during sustained cytotoxic engagement is markedly asymmetric. GzmB recovers after its initial depletion in a protein-synthesis-dependent manner, whereas granzyme A, cathepsin D, serglycin and, most strikingly, mature perforin do not show comparable restoration over the same time window. The distinction between total and mature perforin is important in this context. Using antibodies that discriminate these pools ^14,27^, we detected residual total perforin while the mature, granule-associated form remained strongly reduced. Thus, our data do not imply that perforin synthesis ceases during prolonged activation. Rather, they indicate that the detectable mature perforin pool is not rebuilt with the kinetics observed for GzmB. This kinetic asymmetry is biologically plausible because perforin availability depends not only on transcription and translation, but also on ER export, Golgi processing, adaptor-mediated vesicular sorting and late proteolytic maturation within secretory lysosomes^28–30^. By contrast, GzmB follows a distinct biosynthetic and trafficking program involving mannose-6-phosphate-dependent targeting and proteolytic maturation within the endolysosomal pathway. Sustained CTL function therefore cannot be understood as simple bulk refilling of the granule compartment; individual components of the cytotoxic machinery follow distinct regenerative trajectories. Recent syntheses similarly emphasize that granzymes can act in both canonical, perforin-dependent and debated non-canonical settings, underlining the need to describe how effector composition changes over time^31,32^.

This molecular asymmetry has functional consequences. GzmB recovery was not merely an intracellular accumulation phenomenon: previously engaged CTLs remained able to transfer enzymatically active GzmB to fresh target cells, induce caspase-3 activation and execute productive killing. Moreover, inhibition of GzmB markedly impaired sustained cytotoxicity, whereas inhibition of FasL or calcineurin-dependent signaling had little detectable effect within the experimental window. Together, these observations link replenishment to redeployment and place GzmB at the center of the sustained response under our experimental conditions. They also distinguish this adaptation from the serial-killing transition described in NK cells, in which progressive granule depletion is accompanied by increasing reliance on death-receptor-mediated cytotoxicity^8^. Human CTLs likewise upregulated FasL and inflammatory cytokines (IFNγ and TNFα) during prolonged engagement, demonstrating that alternative effector pathways were induced. Yet these pathways did not replace GzmB as the major detectable contributor to late killing. Serial cytotoxic adaptation may therefore differ fundamentally between cytotoxic lymphocyte populations: rather than abandoning the granule-associated program, CTLs can selectively preserve one of its principal effectors while the remainder of the canonical machinery is incompletely restored. This interpretation is also consistent with studies of CAR T cells in which Fas/FasL contributes to, but does not necessarily dominate, target-cell elimination^33^.

How GzmB gains productive access to target cells in this remodeled state remains an important unresolved question. Target-cell Ca²⁺ dynamics provide evidence that late killing is qualitatively different from the acute response. Fresh CTLs, containing abundant mature perforin, induced the rapid and repetitive Ca²⁺ elevations characteristic of strong membrane perturbation, consistent with previous work showing that perforin pore formation produces rapid ionic fluxes and initiates membrane-repair responses^15,34,35^. In contrast, targets attacked by pre-exposed CTLs displayed weaker, frequently single or gradually developing Ca²⁺ responses, despite subsequent GzmB activity, caspase activation and death. These observations are consistent with a reduced contribution of rapid, overt pore-mediated membrane injury, but they do not establish perforin-independent GzmB delivery. Residual mature perforin below our detection threshold could remain sufficient for GzmB entry. Alternatively, GzmB may be internalized before cytosolic release, with small amounts of perforin acting at intracellular membranes, as proposed in endosomal models of granzyme delivery^35^. Supramolecular attack particles (SMAPs), are reported to kill with delayed kinetics compared with lytic granules, and may provide an additional precedent for extracellular structures enriched in cytotoxic effectors, including GzmB ^36–39^, although our data do not establish that the GzmB-containing compartments identified here generate, or are equivalent to, SMAPs. Resolving these possibilities will require direct manipulation of perforin during the sustained phase and definition of the fate of newly synthesized GzmB.

The intracellular organization of GzmB also changes as cytotoxicity proceeds. GzmB and perforin initially occupy largely overlapping vesicular structures, but progressively become spatially uncoupled, with an increasing proportion of GzmB localized to compartments containing CD107a, CD63 or CD81 but little detectable mature perforin. Expression of these vesicular markers was itself sensitive to inhibition of protein synthesis, indicating that prolonged activation is accompanied by active molecular renewal of the vesicular machinery. This remodeling occurs in the context of known granule-recycling pathways. Following exocytosis, cytotoxic-granule membrane components can be endocytically retrieved and routed through endosomal compartments, and recycled granules can subsequently be reloaded with GzmB before being redeployed to new immunological synapses^40^. These observations argue that GzmB replenishment is coupled to intracellular reorganization rather than simple restoration of the pre-existing granule architecture. Whether the resulting GzmB-containing compartments represent intermediates in lytic-granule re-biogenesis, recycled granule membranes undergoing cargo reloading, alternative secretory organelles, or a mixture of these states remains unresolved. Nevertheless, the molecular and spatial data converge on the same conclusion: the cytotoxic apparatus present after several hours of target-cell engagement is compositionally distinct from that available at the beginning of the response.

A second consequence of sustained engagement is a broadening of cytotoxic behavior. At baseline, CTLs displayed a relatively narrow distribution of intracellular GzmB, whereas prolonged activation produced a substantially broader distribution of GzmB abundance. In parallel, single-cell imaging revealed increased variability in the time required to kill individual target cells, with pre-exposed CTLs displaying a broader range of cytotoxic kinetics and, on average, slower target-cell death. Intraclonal heterogeneity in CTL killing capacity has previously been documented during sustained cytotoxic responses^12^ and can strongly influence the efficiency of tumor-cell elimination in vivo^16^. Our observations suggest that such heterogeneity should not necessarily be viewed as a static property that precedes activation. Sustained engagement itself is accompanied by progressive diversification of effector content and functional kinetics. Whether differences in GzmB renewal directly determine the subsequent killing capacity of individual CTLs will require longitudinal tracking of molecular state and cytotoxic performance in the same cells, but the parallel emergence of molecular and functional dispersion suggests that cytotoxic competence becomes progressively less uniform as the response proceeds.

The persistence of this remodeling during interaction with resistant melanoma cells further indicates that it is not restricted to highly permissive target-cell conditions. Melanoma cells provide a stringent context in which to examine sustained cytotoxicity because they can actively counter CTL attack at the immunological synapse. We previously showed that melanoma cells deploy a lysosomal secretory response capable of limiting perforin-mediated cytotoxicity^14^, and subsequently identified Ca²⁺-dependent synaptic defense programs triggered by CTL attack^15^. Despite this resistance, CTLs engaging melanoma targets followed a global temporal remodeling broadly similar to that observed with sensitive B-cell targets. At the sustained stage, however, melanoma engagement was associated with still greater GzmB abundance and heterogeneity while mature perforin remained low. Thus, target-cell resistance does not appear to divert CTLs toward an entirely different program within the time frame examined; instead, it coexists with, and may intensify, features of the same GzmB-biased remodeling. One possibility is that prolonged or repeated unsuccessful encounters reinforce effector renewal, although establishing the causal basis of this amplification will require additional experiments. This observation is particularly relevant in light of in vivo work showing that resistant tumor cells can require temporally accumulated cytotoxic hits for elimination^19^. In that study, sublethal damage inflicted by individual CTLs could be repaired by the tumor cell unless successive hits arrived closely enough in time, so that the outcome depended as much on target-cell recovery dynamics as on CTL-intrinsic function. Under such conditions, maintaining access to GzmB after the initial granule pool has been deployed could help preserve cytotoxic pressure even when individual killing events become slower or less efficient.

Several limitations define the boundaries of this model. Most mechanistic experiments were performed using highly differentiated human CTL clones and, for detailed phenotyping, under saturating antigenic stimulation to synchronize target recognition. Importantly, however, the core sequence of acute effector loss followed by selective GzmB recovery was also detected at 10 nM peptide, a condition associated with much more limited cytokine production^10,11^, arguing against the remodeling being simply a consequence of maximal inflammatory activation. Whether the same quantitative hierarchy of effector renewal occurs in polyclonal tumor-reactive CTLs, tissue-resident populations or therapeutic T cells remains to be established. Likewise, the transcriptional, translational and trafficking mechanisms that preferentially restore GzmB remain unknown, as does the extent to which antigen strength, repeated synaptic encounters and signals from the tissue microenvironment modulate this process. Finally, we cannot formally exclude a contribution of perforin remaining below our detection threshold, nor distinguish between the possible non-canonical routes of GzmB entry discussed above.

Together, our findings support a two-stage model of CTL cytotoxicity. The acute response draws predominantly on a preassembled perforin-granzyme arsenal capable of rapid membrane perturbation and target-cell elimination. Early serial killing can be supported by conservation and recycling of this pre-existing machinery, but continued engagement ultimately requires active re-arming. Our data indicate that this re-arming is itself selective rather than uniform. Continued engagement does not simply restore this starting configuration. Instead, the cytotoxic machinery is selectively rebuilt around GzmB, which is resynthesized, redistributed and redeployed while the detectable mature perforin pool remains limited. CTLs thereby retain the capacity to kill newly encountered targets, but the mode and kinetics of cytotoxicity become progressively altered and heterogeneous. Sustained killing should therefore be viewed not as repetition of the initial cytotoxic event, but as a dynamic adaptation of the effector machinery to its own prior use.

This view also suggests a practical direction: strategies that stabilize or exploit the GzmB-centric program could help maintain the activity of therapeutic T cells, including CAR T cells, across repeated tumor encounters^31^.

## Acknowledgements

We thank the flow cytometry core facilities of the Centre de Recherches en Cancérologie de Toulouse (CRCT), especially Manon Farcé for technical support. Part of flow cytometry experiments were performed at the Univ Toulouse, INSERM, CNRS, Infinity core facility connected to Toulouse Réseau Imagerie network, member of the France-BioImaging national infrastructure supported by the French National Research Agency (ANR-24-INBS-0005 FBI BIOGEN). We thank Fatima l’faqihi for her expert advice on the experimental set up. This work was supported by the Fondation ARC pour la recherche sur le cancer, and an IUCT-Oncopole Translational Research Grant secured by R.K. And partially supported by the European Research Council (ERC) under the European Union’s Horizon 2020 Research and Innovation Program (Grant agreement No. Syn-951329) secured by S.V. The funders had no role in study design, data collection and interpretation, or preparation of the manuscript.

## Author contributions

R.K. conceived and led the study, designed the project, supervised the research, analyzed and interpreted the data, secured funding, and wrote the manuscript. H.R.G. contributed to study design, performed the imaging and flow-cytometry experiments, analyzed the data, and contributed to manuscript preparation. A.L.I. developed the flow-cytometry panel and analysis pipeline. A.M. prepared the figures. S.M. generated the CTL clones and established the cellular model. S.V. contributed to discussing results and manuscript revision. All authors reviewed and approved the final manuscript. The authors declare no competing interests.

## Material and methods

### Cell models

HLA-A2-restricted CD8^+^ T cells specific for the NLVPMVATV peptide of the cytomegalovirus protein pp65 (CMV specific) clones already available in the lab are used in this study. Cells are cultured in RPMI 1640 medium supplemented with 5% human AB serum (Inst. Biotechnologies J.BOY), 50µmol/L 2-mercaptoethanol, 10mM HEPES, 1X MEM NEAA (Gibco), 1X Sodium pyruvate (Sigma), 10µg/ml ciprofloxacine (AppliChem), 100 IU/ml human rIL-2 and 50 ng/ml human rIL-15 (Miltenyi). T cell clones are stimulated in complete RPMI/HS medium containing 1 mg/ml PHA with 1 x 10^6^ per ml 35 Gy irradiated allogeneic peripheral blood mononuclear cells (isolated on FicollPaque Gradient from blood of healthy donors). Re-stimulation of clones is performed every 2 weeks.

As target cells, EBV-transformed B cells (JY) HLA-A2^+^ are used with high sensitivity to CTL attack that are routinely used in our lab to study CTL behavior. D10 cells (isolated from metastatic melanoma patients, kindly provided by Dr G. Spagnoli, Basel, Switzerland). Target cells are cultured in RPMI 1640 GlutaMAX supplemented with 10% FCS and 50µmol/L 2-mercaptoethanol, 10mM HEPES, 1X MEM NEAA (Gibco), 1X Sodium pyruvate (Sigma), 10µg/ml ciprofloxacine (AppliChem).

All cell lines are screened biweekly for mycoplasma contamination using MycoAlert mycoplasma detection kit (Lonza).

### CTL/Target cell conjugation

Target cells were left unpulsed or pulsed with indicated concentration of antigenic peptide (pp65: NLVPMVATV) during 2h at 37°C/5% CO2, washed three times, re-suspended in RPMI 5% FCS/HEPES (R5). CTLs are pre-stained with 0.1µM CMFDA (5-chloromethylfluorescein diacetate (Molecular Probes)), or CTV (Celltrace Violet (Molecular Probes)) for 15min at 37°C/5% CO2, washed, re-suspended in R5 and added to the target cells at different CTL/target cell (E:T) ratio. For flow cytometry based read-outs, cells are seeded to a 96-wells U-bottom plate and pelleted for 1min, 1500rpm and subsequently incubated at 37°C/5% CO2 for the indicated time point. For imaging based analysis, CTL/target cells seeding is performed either in 15-well chambered slides (Ibidi, Munich, Germany) or in slides.

### CTL cytotoxicity assay

CTL/target cell conjugates are transferred on ice and disrupted by several rounds of pipetting. Before reading samples on flow-cytometry, 0.25 μg of 7-aminoactinomycin D (7-AAD; Waters Biosciences) is added to each sample to measure target-cell death. Alternatively, target cells were pre-loaded with a caspase-3 substrate (NucView® 488, Biotium) at 5 µM in complete medium for 30 min at 37 °C, washed twice, and conjugated with CTLs for the indicated time points. Cytotoxicity was read as the frequency of NucView-positive target cells (excitation 488 nm, emission ∼530/30). In some experiments cytotoxicity was assessed in the presence of Granzyme B inhibitor (200µM z-AAD-CMK, Sigma)

### Functional phenotyping of CTL with flow-cytometry

Using the same experimental set up, CTL/target cell conjugates were stopped at different time points and subjected to staining for the expression of several markers involved in CTL activation and degranulation. CTL/target cells conjugates are washed in ice-cold PBS containing 1% human serum, 1% FBS and 0.5 mM EDTA. Non-specific binding is prevented by 15 min incubation with PBS containing 1% FCS which is used throughout the procedure as staining and washing buffer. Cells were stained first extracellularly with a mix of specific antibodies (anti-CD3 and anti-CD107a) for 30 minutes on ice. Samples are washed thoroughly and fixed with 3% PFA, permeabilized with 0.1% saponin (in PBS/3% BSA/HEPES), and stained for intracellular markers (anti-perforin and anti-GranzymeB). Samples were acquired using LSR II (Becton and Dickinson) or Symphony A5 (Waters Biosciences). Results are analyzed using the FlowJo Pro software (Tree Star, Inc.). Where indicated, CTL/target cell conjugation was performed in the presence of 1 µg/ml Cyclosporin A (CsA) or 10 µg/ml cycloheximide/CHX for blocking NFAT signaling or protein de novo synthesis respectively. The following mouse anti human antibodies were used: Alexa Fluor APC-Cy7 or Alexa Fluor 647 anti-perforin (5 μg/ml, clone dG9; BioLegend), BV421 anti-perforin (5 μg/ml, clone B-D48; BioLegend), BUV395 anti-CD107a (10 μg/ml, clone H4A3; Waters Biosciences), Pacific Blue anti-granzyme B (10 μg/ml, clone GB11; BioLegend) and PE anti-CD3 (10 μg/ml, clone OKT3; Waters Biosciences), BV783 anti-CD3 (10 μg/ml, clone SK7; Waters Biosciences), BV570 anti-CD8 (10 μg/ml, clone RPA-TB; Waters Biosciences), BV711 anti-CD69 (10 μg/ml, clone FN50; Waters Biosciences), Alexa Fluor 594 anti-Cathepsin B (10 μg/ml, clone H.5; Santa Cruz), Alexa Fluor 680 anti-Rab27a (10 μg/ml, clone E.8; SantaCruz), RB744 anti-TNFα (10 μg/ml, clone N297A; Waters Biosciences), APC anti-IFNy (10 μg/ml, clone 4S,B3; Waters Biosciences), BB700 anti-IL-2 (10 μg/ml, clone MQ1-17H12; Waters Biosciences), PE-Fire700 anti-PD1 (10 μg/ml, clone A17188B; BioLegend), PE-Cy7 anti-LAG-3 (10 μg/ml, clone 11C3C65; Waters Biosciences), SuperBright783 anti-TIM3 (10 μg/ml, clone Ber-ACT8; Waters Biosciences).

### High dimensional analysis

CTLs were identified as live CMFDA⁺ cells, and were exported as FCS files and analyzed using FlowJo v10 with the Opt-SNE and FlowSOM plugins. Before dimensionality reduction, fluorescence intensities were transformed using an arcsinh transformation with a cofactor of 150. Equal numbers of CTLs were randomly downsampled from each condition and concatenated to avoid bias due to unequal cell numbers. The markers used for dimensionality reduction and clustering included CD3, CD107a, GzmB, GzmA, perforin, FasL, IFNγ, TNFα, cathepsin D, serglycin, CD25, CD69, PD-1, TIM-3, LAG-3. Markers used only for gating, cell identification, or viability exclusion were not included in the Opt-SNE or FlowSOM computation. Opt-SNE analysis was performed on the concatenated CTL population using 2000 events per condition, with the following parameters: perplexity 30, iterations 1000, automatic learning rate, and theta 0.5. The resulting Opt-SNE maps were visualized according to time point, antigenic stimulation, target-cell type, or FlowSOM cluster assignment, as indicated in the corresponding figure legends. FlowSOM clustering was performed on the same transformed and downsampled CTL dataset using the selected phenotypic markers. The self-organizing map was generated using a 10 × 10 grid, followed by metaclustering into 7 phenotypic clusters. Cluster identity was assigned based on marker-expression profiles and visualized on the corresponding Opt-SNE maps. Cluster frequencies were calculated as the percentage of live CTLs within each condition. Median fluorescence intensity values for individual markers were extracted from manually gated CTLs or from FlowSOM-defined clusters, as indicated. For pooled quantifications, values were normalized to the corresponding non-pulsed condition or to the indicated baseline condition within each independent experiment before statistical analysis.

### Confocal imaging

Target cells either unpulsed or pulsed with 10 µM of antigenic peptide and conjugated for indicated time duration with CTLs. Conjugates were stopped by washing in ice-cold PBS containing 1% human serum, 1% FBS and 0.5 mM EDTA. Cells are fixed with 3% paraformaldehyde (PFA) for 10min at room temperature, washed and permeabilized with 0.1% saponin (in PBS/3% BSA/HEPES), and stained intracellularly for perforin, GzmB and CD107a for 1 hour at room temperature. The samples were mounted in 90% glycerol-PBS containing 2.5% DABCO (Fluka) and examined using LSM 780 (Zeiss) confocal microscope over a x63 Plan-Apochromat objective (1.4 oil). Electronic zoom 4. Z-Stack images of optical sections were acquired throughout the cell volume. Images are scored by evaluating, for each experimental condition, at least 20 CTL/target cell conjugates in randomly selected fields from at least three independent experiments. For co-localization analysis, 3D image stacks were processed using Fiji (ImageJ) and Imaris (Bitplane) software. Co-localization of perforin and/or GzmB was assessed using voxel-based overlap and intensity correlation analysis using Imaris software. Pearson’s correlation coefficients were calculated for each channel pair. Quantification was performed on at least 30 individual CTLs per condition, across three independent experiments. Representative images were selected for display, and all settings were kept constant between conditions for comparative analysis.

### Live cell imaging

Target cells preloaded with 10µM of antigenic peptide are seeded at 1.5 × 10^5^ cells per well on poly-D-lysine-coated 15-well chambered slides 5 min before imaging. Chambered slides were mounted on a heated stage within a temperature-controlled chamber maintained at 37 °C, and constant CO_2_ concentrations (5%). To monitor intracellular calcium ([Ca²⁺]i) dynamics, target cells were preloaded with Fluo-4 AM (2 μM, Molecular Probes) in imaging buffer for 30 min at 37 °C, washed twice, and kept in phenol red-free medium until acquisition. Effector CTLs (2 × 10⁵ cells per well), either fresh or previously conjugated with targets, were added at the start of recording. CTLs were optionally labeled with 1 μM Cell Tracer Violet (Invitrogen) for visualization. To simultaneously monitor membrane permeabilization, propidium iodide (PI; 0.5 μg/mL, Sigma) was included in the medium. A rise in PI fluorescence was interpreted as a marker of membrane disruption, often associated with perforin pore formation. Images of CTL/target cell conjugates were acquired for 1 h using either a Nikon spinning disk microscope (60×/1.4 NA Plan Apochromat LBDA objective lens (Nikon Instruments)). Image acquisition was controlled using MetaMorph Software (Version 7.10.5.476, Molecular Devices) and Modular V2.0 GATACA software. A sCMOS Hamamatsu ORCA-Flash4.0 V3 camera or an emCCD Evolve (Photometrics) camera are used for acquisitions. Target cells were preloaded with 5 ng/µl Fluo-4 AM, (Molecular Probes) during 30 min and washed before seeding in chambered slides. Image analysis is performed using Image J software.

Fluo-4 fluorescence intensity was measured in individual target cells using predefined masks based on CMFDA-positive CTLs and Fluo-4 signal, using Fiji (ImageJ) software. For each cell, normalized fluorescence intensity (ΔF/F₀) was calculated, and calcium flux kinetics were plotted over time. Single-cell calcium response types were categorized as sharp, repetitive spikes, gradual increases, or absent responses. At least, 40 conjugates per condition were analyzed across three independent experiments. PI internalization time and final intensity was measured using ImageJ software.

### Granzyme B activity assay

To monitor granzyme B (GzmB) activity specifically delivered to target cells by CTLs, the GranToxiLux assay kit (OncoImmunin, GTLX-488) was used according to the manufacturer’s instructions. Briefly, target cells were preloaded with the fluorogenic GzmB substrate provided in the kit (a cell-permeable peptide conjugated to a fluorescent reporter), which becomes fluorescent upon cleavage by active GzmB. After labeling, targets were washed and conjugated with CTLs for indicated time intervals. Granzyme B activity in target cells was then assessed by flow cytometry as the percentage of GranToxiLux⁺ cells.

## References

1. McKenzie, B., Khazen, R. & Valitutti, S. Greek Fire, Poison Arrows, and Scorpion Bombs: How Tumor Cells Defend Against the Siege Weapons of Cytotoxic T Lymphocytes. Front. Immunol. 13, (2022).

2. Romero, P., Cerottini, J. C. & Speiser, D. E. The Human T Cell Response to Melanoma Antigens. Advances in Immunology vol. 92 Preprint at 10.1016/S0065-2776(06)92005-7 (2006).

3. Romero, P. et al. Ex Vivo Staining of Metastatic Lymph Nodes by Class I Major Histocompatibility Complex Tetramers Reveals High Numbers of Antigen-experienced Tumor-specific Cytolytic T Lymphocytes. J. Exp. Med. 188, 1641–1650 (1998).

4. Tumeh, P. C. et al. PD-1 blockade induces responses by inhibiting adaptive immune resistance. Nature 515, 568–571 (2014).

5. Huang, A. C. & Zappasodi, R. A decade of checkpoint blockade immunotherapy in melanoma: understanding the molecular basis for immune sensitivity and resistance. Nature Immunology vol. 23 Preprint at 10.1038/s41590-022-01141-1 (2022).

6. Sade-Feldman, M. et al. Defining T Cell States Associated with Response to Checkpoint Immunotherapy in Melanoma. Cell 176, 404 (2019).

7. McKenzie, M. D. et al. Perforin and Fas induced by IFNγ and TNFα mediate beta cell death by OT-I CTL. Int. Immunol. 18, (2006).

8. Prager, I. et al. NK cells switch from granzyme B to death receptor–mediated cytotoxicity during serial killing. Journal of Experimental Medicine 216, 2113–2127 (2019).

9. He, J.-S. & Ostergaard, H. L. CTLs Contain and Use Intracellular Stores of FasL Distinct from Cytolytic Granules. The Journal of Immunology 179, 2339–2348 (2007).

10. Wiedemann, A., Depoil, D., Faroudi, M. & Valitutti, S. Cytotoxic T lymphocytes kill multiple targets simultaneously via spatiotemporal uncoupling of lytic and stimulatory synapses. Proc. Natl. Acad. Sci. U. S. A. 103, (2006).

11. Faroudi, M. et al. Lytic versus stimulatory synapse in cytotoxic T lymphocyte/target cell interaction: Manifestation of a dual activation threshold. Proceedings of the National Academy of Sciences 100, 14145–14150 (2003).

12. Vasconcelos, Z. et al. Individual Human Cytotoxic T Lymphocytes Exhibit Intraclonal Heterogeneity during Sustained Killing. Cell Rep. 11, (2015).

13. Caramalho, Í., Faroudi, M., Padovan, E., Müller, S. & Valitutti, S. Visualizing CTL/melanoma cell interactions: multiple hits must be delivered for tumour cell annihilation. J. Cell. Mol. Med. 13, 3834–3846 (2009).

14. Khazen, R. et al. Melanoma cell lysosome secretory burst neutralizes the CTL-mediated cytotoxicity at the lytic synapse. Nat. Commun. 7, (2016).

15. Filali, L. et al. Ultrarapid lytic granule release from CTLs activates Ca2+-dependent synaptic resistance pathways in melanoma cells. Sci. Adv. 8, (2022).

16. Khazen, R. et al. Functional heterogeneity of cytotoxic T cells and tumor resistance to cytotoxic hits limit anti-tumor activity in vivo. EMBO J. 40, (2021).

17. Khazen, R., Müller, S., Lafouresse, F., Valitutti, S. & Cussat-Blanc, S. Sequential adjustment of cytotoxic T lymphocyte densities improves efficacy in controlling tumor growth. Sci. Rep. 9, (2019).

18. Halle, S. et al. In Vivo Killing Capacity of Cytotoxic T Cells Is Limited and Involves Dynamic Interactions and T Cell Cooperativity. Immunity 44, (2016).

19. Weigelin, B. et al. Cytotoxic T cells are able to efficiently eliminate cancer cells by additive cytotoxicity. Nat. Commun. 12, (2021).

20. Halle, S., Halle, O. & Förster, R. Mechanisms and Dynamics of T Cell-Mediated Cytotoxicity In Vivo. Trends Immunol. 38, 432–443 (2017).

21. Bertrand, F. et al. An initial and rapid step of lytic granule secretion precedes microtubule organizing center polarization at the cytotoxic T lymphocyte/target cell synapse. Proceedings of the National Academy of Sciences 110, 6073–6078 (2013).

22. Ming, M., Schirra, C., Becherer, U., Stevens, D. R. & Rettig, J. Behavior and Properties of Mature Lytic Granules at the Immunological Synapse of Human Cytotoxic T Lymphocytes. PLoS One 10, e0135994 (2015).

23. Sanchez-Ruiz, Y., Valitutti, S. & Dupre, L. Stepwise Maturation of Lytic Granules during Differentiation and Activation of Human CD8+ T Lymphocytes. PLoS One 6, e27057 (2011).

24. Isaaz, S., Baetz, K., Olsen, K., Podack, E. & Griffiths, G. M. Serial killing by cytotoxic T lymphocytes: T cell receptor triggers degranulation, re-filling of the lytic granules and secretion of lytic proteins via a non-granule pathway. Eur. J. Immunol. 25, 1071–1079 (1995).

25. Lisci, M. et al. Mitochondrial translation is required for sustained killing by cytotoxic T cells. Science (1979). 374, (2021).

26. Makedonas, G. et al. Rapid Up-Regulation and Granule-Independent Transport of Perforin to the Immunological Synapse Define a Novel Mechanism of Antigen-Specific CD8+ T Cell Cytotoxic Activity. The Journal of Immunology 182, 5560–5569 (2009).

27. Hersperger, A. R., Makedonas, G. & Betts, M. R. Flow cytometric detection of perforin upregulation in human CD8 T cells. Cytometry Part A 73A, 1050–1057 (2008).

28. Brennan, A. J. et al. Protection from Endogenous Perforin: Glycans and the C Terminus Regulate Exocytic Trafficking in Cytotoxic Lymphocytes. Immunity 34, 879–892 (2011).

29. Konjar, Š. et al. Human and mouse perforin are processed in part through cleavage by the lysosomal cysteine proteinase cathepsin L. Immunology 131, 257–267 (2010).

30. Krzewski, K., Gil-Krzewska, A., Nguyen, V., Peruzzi, G. & Coligan, J. E. LAMP1/CD107a is required for efficient perforin delivery to lytic granules and NK-cell cytotoxicity. Blood 121, 4672–4683 (2013).

31. Thompson, R. & Cao, X. Reassessing granzyme B: unveiling perforin-independent versatility in immune responses and therapeutic potentials. Front. Immunol. 15, (2024).

32. McKenzie, B. et al. Perforin-mediated pore formation at the lytic synapse triggers the canonical pyroptotic cell death pathway. Preprint at 10.1101/2024.05.24.595698 (2024).

33. Montalvo, M. J. et al. Decoding the mechanisms of chimeric antigen receptor (CAR) T cell-mediated killing of tumors: insights from granzyme and Fas inhibition. Cell Death Dis. 15, 109 (2024).

34. Lopez, J. A. et al. Perforin forms transient pores on the target cell plasma membrane to facilitate rapid access of granzymes during killer cell attack. Blood 121, 2659–2668 (2013).

35. Thiery, J. et al. Perforin pores in the endosomal membrane trigger the release of endocytosed granzyme B into the cytosol of target cells. Nat. Immunol. 12, 770–777 (2011).

36. Bálint et al. Supramolecular attack particles are autonomous killing entities released from cytotoxic T cells. Science (1979). 368, (2020).

37. Chang, H. F. et al. Identification of distinct cytotoxic granules as the origin of supramolecular attack particles in T lymphocytes. Nat. Commun. 13, (2022).

38. Cassioli, C. & Baldari, C. T. The Expanding Arsenal of Cytotoxic T Cells. Front. Immunol. 13, (2022).

39. Cassioli, C. et al. Activation-induced thrombospondin-4 works with thrombospondin-1 to build cytotoxic supramolecular attack particles. Proceedings of the National Academy of Sciences 122, (2025).

40. Chang, H.-F. et al. Endocytosis of Cytotoxic Granules Is Essential for Multiple Killing of Target Cells by T Lymphocytes. The Journal of Immunology 197, 2473–2484 (2016).

