## Supplementary information for "Selective granzyme B replenishment dominates the molecular switch for sustained cytotoxicity in effector CD8^+^ T lymphocytes"

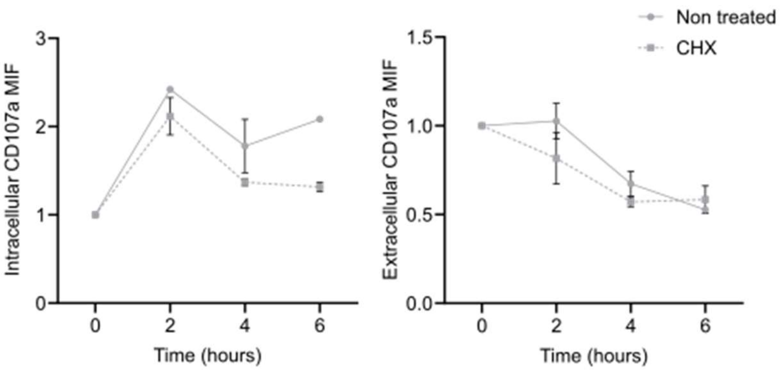


**Supplementary Figure 1**
**Rodriguez et al.**

**Supplementary figure 1.** Time kinetic expression of intra- versus extra-cellular expression of CD107a molecule in CTLs conjugated with peptide pulse target cells. MFI are represented as normalized to non-pulsed condition. Data are shown as mean +/- s.e.m.

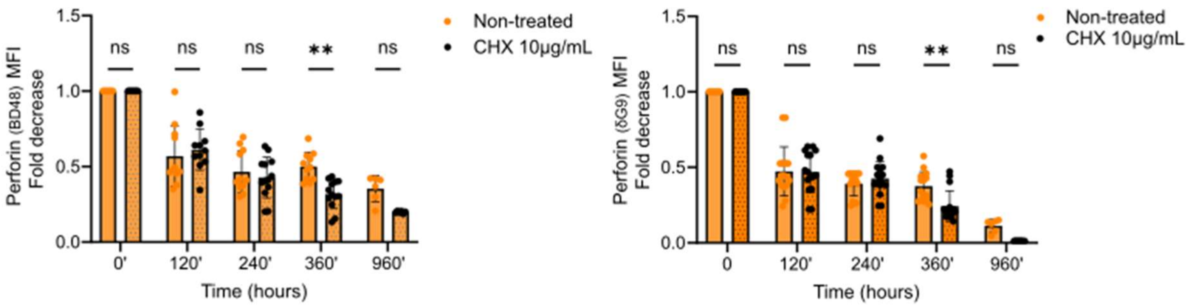


**Supplementary Figure 2**
**Rodriguez et al.**

**Supplementary figure 2.** Time kinetic expression of total perforin expression using clone B-D48 (total perforin) or clone dG9 (mature perforin) antibody in CTLs conjugated with peptide pulse target cells. MFI are represented as normalized to non-pulsed condition. Data are shown as mean +/- s.e.m. Statistical significance was assessed using unpaired t-test. ns, not significant.

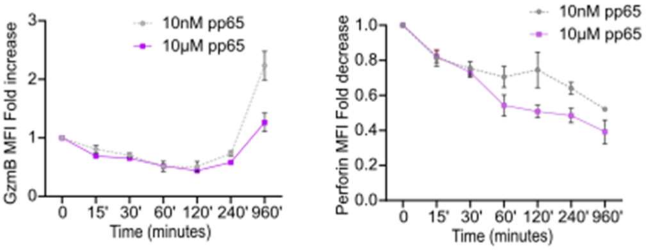


**Supplementary Figure 3**
**Rodriguez et al.**

**Supplementary figure 3.** Time kinetic expression of GzmB and mature perforin expression using clone in CTLs conjugated with target cells either pulsed with 10nM or 10 µM of antigenic peptide. MFI are represented as normalized to non-pulsed condition. Data are shown as mean +/- s.e.m.

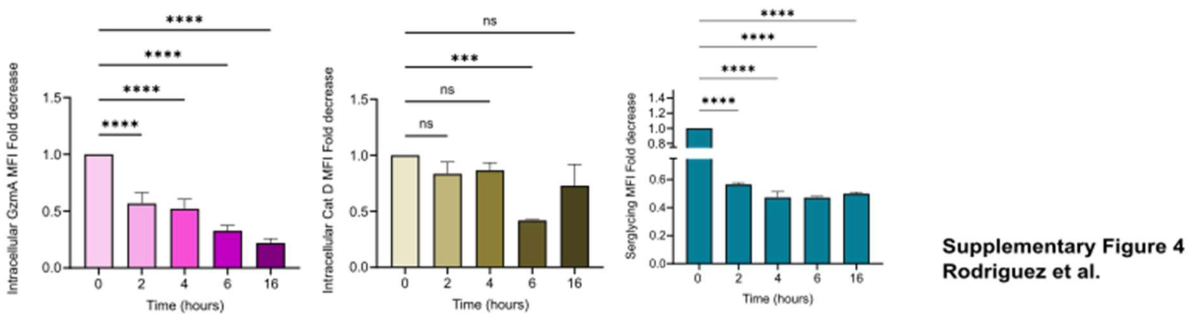


**Supplementary figure 4. Relative expression patterns of key lytic related molecules across time.** Bar plots illustrating time kinetics of relative expression of indicate the proteins in CTLs conjugated with peptide pulsed target cells. MFI are represented as normalized to non-pulsed condition. Data are shown as mean +/- s.e.m. Statistical significance was assessed using unpaired t-test. ns, not significant. *P < 0.05; ***P < 0.001, ****P < 0.0001.

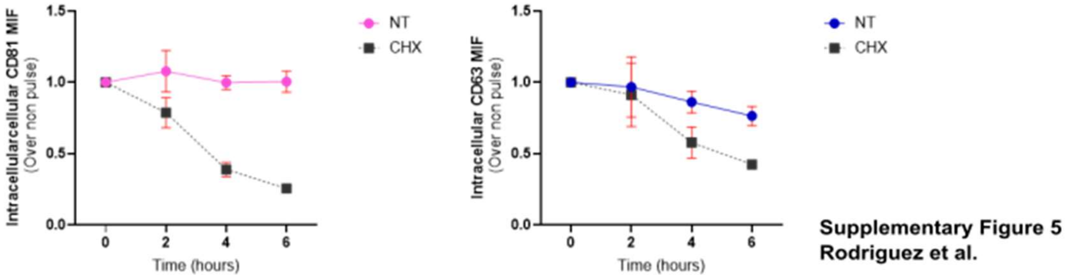


**Supplementary figure 5.** Time kinetic expression of intra-cellular expression of CD63 and CD81 molecules in CTLs conjugated with peptide-pulse target cells. MFI are represented as normalized to non-pulsed condition. Data are shown as mean +/- s.e.m.

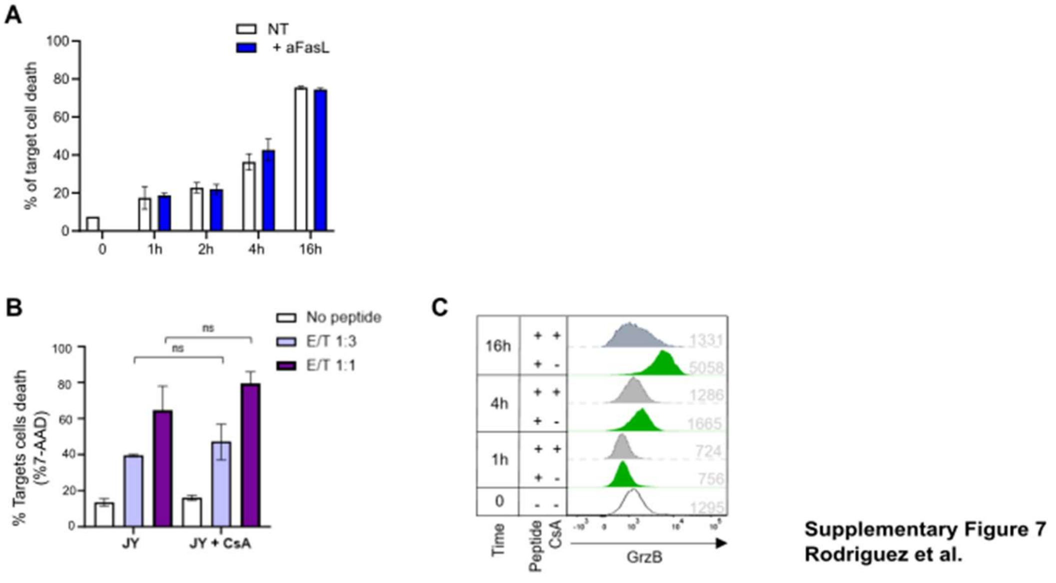


**Supplementary figure 6. Inhibition of FasL or cytokine production does not impair granzyme B replenishment or sustained cytotoxicity.** CTLs were conjugated with target cells either non-pulsed or pulsed with cognate peptide in the presence or absence of cyclosporin A (CsA) or inhibitory FasL antibody. **(A-B)** CTL cytotoxicity quantified as the percentage of 7-AAD$^{+}$ target cells in the presence of FasL inhibitor (A) or CsA (B). **(C)** Representative flow cytometry histograms showing intracellular GzmB expression over time. Data are shown as mean +/- s.e.m. Statistical significance was assessed using unpaired t-test. ns, not significant.

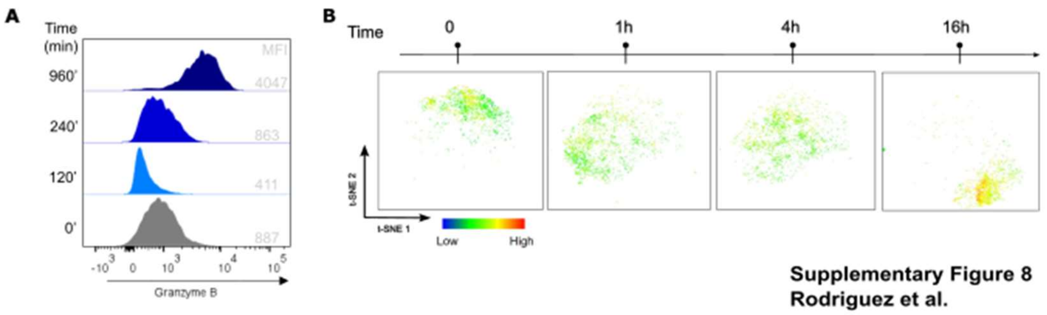


**Supplementary figure 7. Granzyme B replenishment during sustained phase of cytotoxicity.**
CTLs were conjugated with target cells pulsed with cognate peptide for indicated time. Granzyme B content was quantified using flow cytometry. **(A)** Representative histograms showing the distribution of GzmB expression over time. **(B)** Opt-SNE projection of CTLs colored according to per-cell GzmB intensity (pseudocolor scale; each dot represents a single CTL).
